# Temporary pond loss as a result of pasture abandonment: exploring the social-ecological drivers and consequences on amphibians

**DOI:** 10.1101/751248

**Authors:** Nándor Erős, Cristian Malos, Csaba Horváth, Tibor Hartel

**Affiliations:** Hungarian Department of Biology and Ecology and Center of Systems Biology, Biodiversity and Bioresources (Center of ‘3B’), Babes-Bolyai University of Cluj-Napoca, Cluj-Napoca, Romania; Babes-Bolyai University, Faculty of Environmental Science and Engineering, Fântânele street, no. 30, 400294, Cluj-Napoca, Romania

**Keywords:** land abandonment, extensive grazing, amphibian conservation, traditional farming, social capital, Romania, Eastern Europe

## Abstract

Amphibian conservation in farming landscapes should address two challenges. First, to understand the relationship between landuse and amphibian habitat quality and second, to understand and support of the capacity of the local communities to continue those farming practices which supports amphibian friendly habitats. While the first challenge is addressed by several studies, there is virtually no study addressing the socio-economic drivers of landuse change. The major aim of this study to fill this knowledge gap by (i) documenting the temporary pond loss in 10 years in a traditionally managed pasture as a result of land abandonment and (ii) exploring the socio-economic and environmental drivers of abandonment. The results show a dramatic increase of scrub cover in the study area as a result of land abandonment. The formation of temporary ponds was negatively influenced by the increase of scrub cover in the vicinity of ponds. There were no differences between the amphibian species assemblages nor the species richness between the lost- and persisting ponds. The social component of the research highlights possible maladaptive paths in pasture management reinforced by the village depopulation, wrong interpretation of nature protection law by officials, scrub encroachment caused decrease in pasture quality and the demotivation of locals to restart traditional grazing. Conservation efforts in traditional farming landscapes facing land abandonment should (i) target the maximization of the quality of the remaining ponds for amphibians and (ii) should support reviving traditional farming practices within the local community.

## 1. Introduction

Temporary ponds are small natural features with high ecological values (Calhoun et al., 2017; Flitcroft, Boon, Cooperman, Harrison, & Bignoli, 2019). Scattered across the landscape, these ponds increases the local and landscape scale biodiversity for several taxa (Ruhí et al., 2012; Demeter & Hartel, 2007; Scheffer et al., 2006) and contributes to the metapopulation persistence for several species (Semlitsch & Bodie, 1998). The habitat value of small, man-made temporary ponds for amphibians in farming landscapes was documented by several studies (Ruhí et al., 2012; Curado et al., 2011; Hartel and von Wehrden, 2013; Plǎiaşu, Bǎncilǎ, Samoilǎ, Hartel & Cogǎlniceanu, 2012; Buono et al., 2019). In several regions the duration of the temporary ponds is largely dependent on precipitation (Winter, 2000). Temporary ponds can represent proper habitats for amphibians because the periodical desiccation controls predators and allows amphibian larvae to metamorphose (Semlitsch, 2000; Semlitsch, Peterman, Anderson, Drake & Ousterhout, 2015; Semlitsch & Bodie, 1998; Wellborn, Skelly & Werner, 1996). Although some amphibian species are adapted to the risk of pond drying either by accelerating their larval development or by breeding in periods when the water availability is highest (i.e. in spring or in short synchronization with rainfall which fills the ponds) (Wells, 1977; Merilä et al., 2010; Newman, 1992; Richter-Boix, Tejedo & Rezende, 2011), accelerated pond drying can cause reproductive failure (Alford & Richards, 1999; Hartel, Bǎncilǎ & Cogǎlniceanu, 2011).

Farming landscapes can be rich in temporary ponds (Hartel & von Wehrden, 2013) because human activities can result in temporary pond formation. Especially the traditional grazing with cattle can maintain optimal amphibian habitats, especially in Europe (Howell et al., 2019). Cattle grazing may contribute to the maintenance of temporary ponds due to trampling and removal of vegetation which would fill these ponds and/or accelerate their drying (Pyke & Marty, 2005; Winter, 2000). Although the value of traditional grassland management practices (i.e. grazing, mowing) for biodiversity conservation is increasingly recognized in Europe (Bergmeier, Petermann & Schröder, 2010; Halada, Evans, Romão & Petersen, 2011; Plieninger et al., 2015), the continuation of these practices by the local communities faces serious socio-economic challenges (Fischer, Hartel & Kuemmerle, 2012). As a result, many traditionally managed grasslands with high natural values are either intensified (or converted into other landuse types, e.g. crop fields) or completely abandoned (Ustaoglu & Collier, 2018) both having dramatic consequences on temporary ponds as amphibian habitats, resulting in their disappearance or decreasing their duration (Curado et al., 2011; Ferreira & Beja, 2013; Skelly, Werner & Cortwright, 1999).

An effective conservation strategy for amphibians in farming landscapes should address two key interlinked challenges. The first is related to the understanding of the relationships between landuse and the quality of amphibian habitats. The amphibian conservation literature focusing on traditional farming landscapes typically documents this relationship and proposes management interventions based on these results (Buckley, Beebee & Schmidt, 2012; Cogǎlniceanu, Bǎncilǎ, Plǎiaşu, Samoilǎ & Hartel, 2012; Tanadini, Schmidt, Meier, Pellet & Perrin, 2012). The second challenge is related to the understanding and supporting of the capacity of the local communities to continue the amphibian friendly farming practices (Fischer, Hartel & Kuemmerle, 2012). This social aspect is important because the values, socio-economic aspirations and the connections of farmers with their farmland is a key pre-requisite for sustaining high natural and cultural value landscapes and the decline of farmland biodiversity is typically driven by socio-economic changes (Fischer, Hartel & Kuemmerle, 2012; Hartel, Fagerholm, Torralba, Balázsi & Plieninger, 2018; Snoo et al., 2013). In the following we will refer to the simultaneous employment of social and ecological studies to understand and advance amphibian conservation in traditional farming landscapes as ‘holistic’ approaches (Hanspach et al., 2014; Hanspach, Loos, Dorresteijn, Abson & Fisher, 2016).

To the best of our knowledge, holistic approaches for amphibian conservation in traditional farming landscapes are still scarce globally and we are unaware about the existence of such studies in Eastern Europe. To fill this knowledge gap, document the dramatic loss of temporary ponds within a decade, as a result of the abandonment of traditional extensive grazing while providing an understanding on the socio-economic and environmental drivers of land abandonment and amphibian pond loss. Our research is timely because large areas of managed, highly biodiverse grasslands from Eastern Europe face the threat of abandonment with dramatic impact on several species of conservation concern (Ruprecht, 2017). Furthermore the litter accumulation in temporary ponds as a consequence of grazing cessation (Hartel & von Wehrden, 2013), combined with increased evapotranspiration in extreme droughts results in low pond quality and high reproductive failures of amphibians (Pyke & Marty, 2005). The importance of the research is increased by the fact that parts of the studied grassland were also under different forms of nature conservation regulations in the past century, therefore our results can be considered general for similar systems under protected area regulations from Romania and Eastern Europe. Our specific objectives (*O*) were:

*O1.* To describe the persisting and lost temporary ponds after the cessation of the traditional grazing management and the pasture maintenance activities.
*O2.* To explore the changes in scrub cover in the whole study area and relate the persistence of temporary ponds to the scrub cover in their vicinity. We interpreted the increase of scrub cover in the vicinity of the temporary ponds as a proxy for increasing water evaporation and/or increasing plant litter accumulation due to the cessation of grazing and woody vegetation management.
*O3.* To explore the aquatic habitat use by amphibians and the extent at which the pond loss affects their breeding opportunities.
*O4.* To explore the socio-economic and environmental causes and mechanisms of the management and abandonment of traditional grazing and its consequences on temporary ponds. This objective provides a socio-economic background knowledge about the social challenges of temporary pond conservation in traditional rural regions facing dramatic socio-economic changes.

## 2. Methods

### 2.1 Study area

The study area is situated at the periphery of the Transylvanian Plain (46°50′28″N, 23°38′31″ E) and the steppe and continental biogeographic regions of Romania and it is a pasture traditionally managed by cattle, buffalo and sheep grazing, with the associated scrub clearing and drinking trough maintaining activities. Traditionally the pasture management was communal (*sensu* Sutcliffe et al., 2013). Details about the trends of annual temperatures and precipitations in the past two decades will be presented below (Figure 1). A 1.5 ha of the pasture was under protected area regulation (botanical reserve) since 1932 (reinforced by Law no. 5/2000) because of its outstanding plant biodiversity (*cca*. 1400 plant species Soó, 1927).

**Figure 1.**
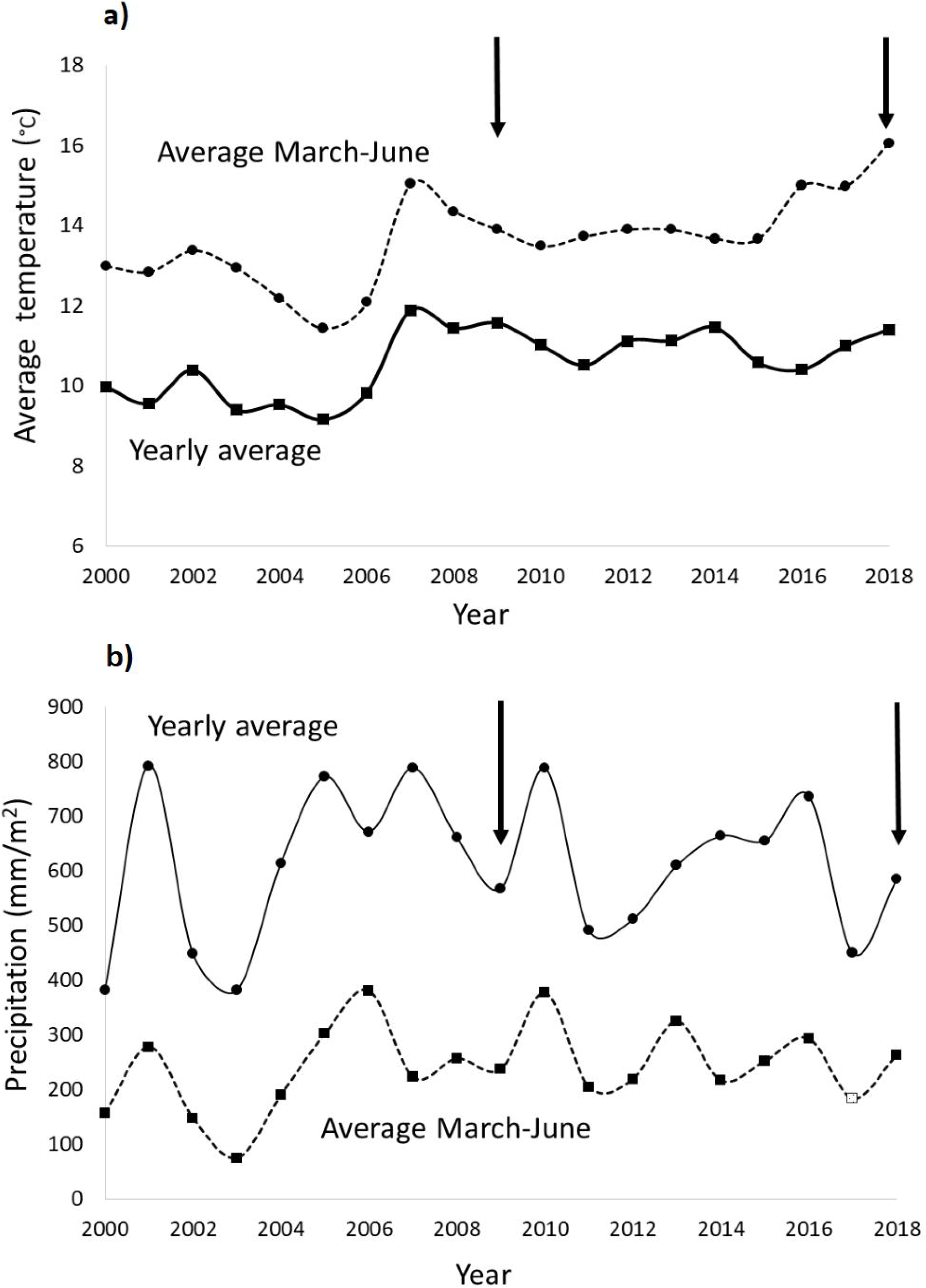
Temperature (a) and precipitation (b) data for the period of 2000-2018. The yearly averages as well as the averages for March-June (i.e. the breeding season and larval periods of amphibians) are shown. The vertical arrows show the pond and amphibian survey years (2019 not shows because the lack of weather data).

The protected status was reinforced in 2004 and the protected area surface extended to 97 ha (Government Decision no. 2151/2004) also covering traditionally managed pastures. Herpetologically important endemic subspecies which were described from the study area are the Transylvanian Smooth Newt (*Lissotriton vulgaris ampelensis*) (Sos & Hegyeli, 2014) and the Hungarian Meadow Viper (*Vipera ursinii rakosiensis*) (first record Bielz, 1888). The studied pasture has only small sized stagnant water bodies. The five ponds with permanent character were documented decades ago (Nyárády, 1941) while the other ponds were temporary and not documented in historical records. The temporary ponds from the study area strongly depend on the amounts of precipitation. The hydroperiod of the temporary ponds from such systems and regions varies in average from 6 to 10 weeks (Hartel, Bǎncilǎ & Cogǎlniceanu, 2011).

### 2.2 Weather conditions

Given that the only source of water for the temporary ponds in the study area is the precipitation, we will present the fluctuation of temperature and precipitation for the period of 2000-2018 (the 2019 data being not available at the period of manuscript development) (ECA&D project dataset, 2019). We present yearly averages as well as the averages of the period of March-June, i.e. the breeding season and larval development period for amphibians from this region. We selected the period of 2000-2018 in order to capture a broader temporal pattern of the weather conditions, which may influence the duration of the temporary ponds (either through evapotranspiration in a given year or by increasing soil drought). In the case of average yearly temperatures, the yearly average values showed a slight, while the spring (March-June) averages showed a more marked increase since 2007 and in 2018 (Figure 1). In the case of the average yearly precipitation the lowest extremes were in 2000, 2002, 2003, 2011, 2014 and 2017 and no trend is apparent in the case of spring precipitation (Figure 1).

### 2.3 Pond and amphibian sampling

Five permanent ponds (ranging from 0.1 ha to 1.2 ha) which were first described in 1941 (Nyárády, 1941) were identified in this study as well. Besides the permanent ponds we carried out a comprehensive temporary pond sampling across the pasture in 2009, 2018 and 2019. The 2009 survey was carried out in the middle and second part of April while in 2018 and 2019 the surveys were conducted between early March and late May, with occasional visits in June. The ponds were located with a GPS device (accuracy <10 m) in all three years.

Amphibians were searched in all three years (2009, 2018 and 2019). Ponds were visited multiple times in all three periods and the surveys captured the periods when the detectability of the different amphibian species based on the presence of adults, eggs or larvae in the ponds were most likely. In 2009 the amphibian surveys were carried out in the middle and second part of April while in 2018 and 2019, each pond was surveyed in March, April and May. At the end of May the temporary ponds started to dry (Figure 2), and occasional visits in June showed that no temporary ponds were formed for more than 10 days, despite the rains. Amphibians were searched with dip-netting but we considered also the visual inspections (in the case of ephemeral ponds) as well as the detection of calling males. The temporary ponds were accessible across their whole surface. In the case of the permanent ponds we sampled five approximatively 15-20 m^2^ areas along the shore of the ponds, where the shoreline was accessible. If we identified any life stage of a species in a ponds, we considered the species present and if no life stage was detected we considered the species absent. In all three years we observed adults, eggs and larvae of most of amphibian species in the majority of the ponds surveyed (Hartel and Erős *unpublished*) therefore we believe that with the information gathered we can infer the breeding habitat loss resulting from the pond loss in the study area (see also Skelly, Werner & Cortwright, 1999).

**Figure 2.**
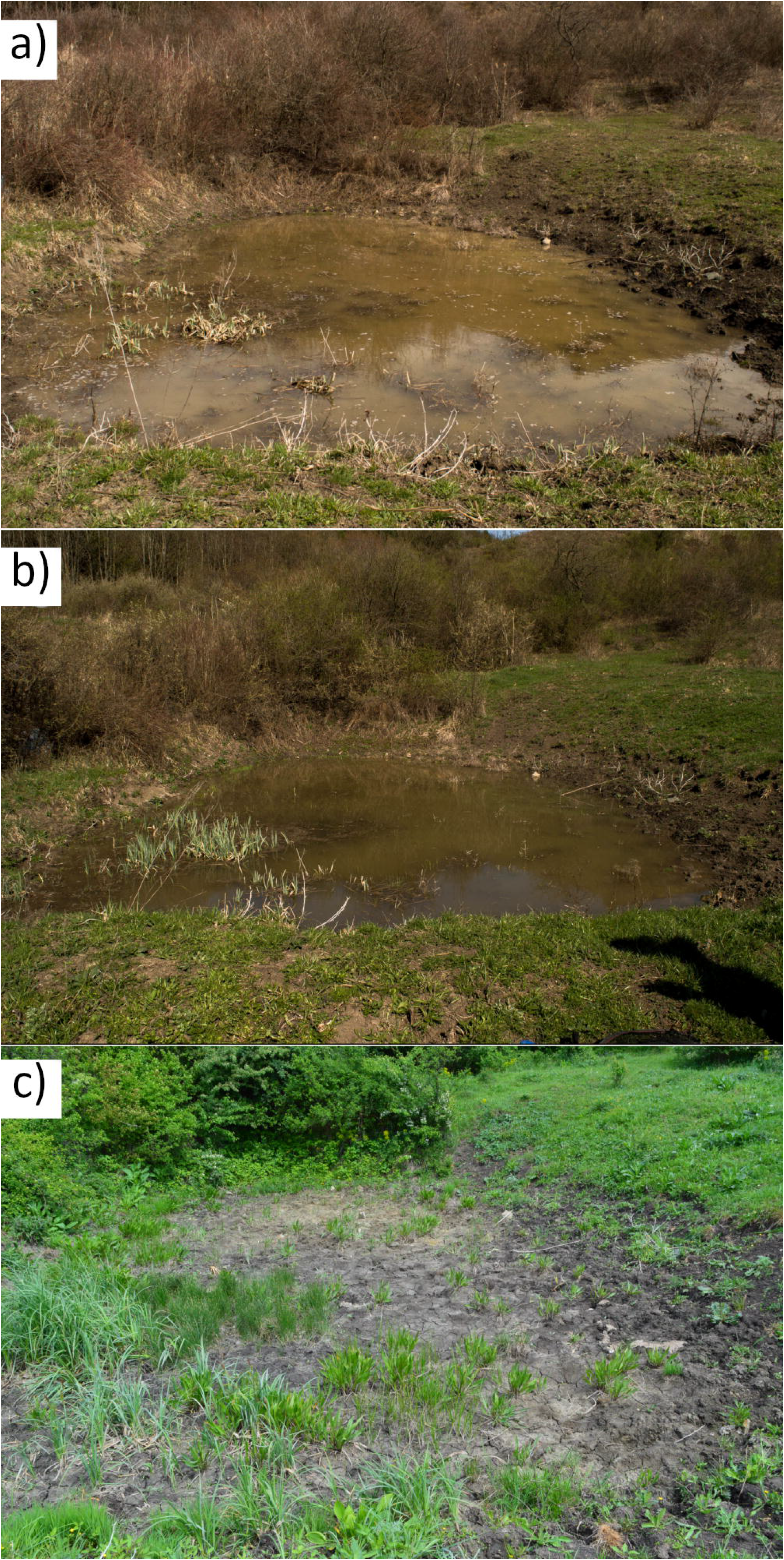
An example of temporary pond characteristic for the study area. The pond was dried till the end of May both in 2018 and 2019. While four amphibian species bred in this pond, none metamorphosed because of pond drying. The picture series represent a pond in end of March (a), April (b) and early May (c) period.

### 2.4 Pond and scrub variables

For each temporary pond we recorded the following three variables: pond area (visually estimated in m^2^), the maximum depth (cm), scrub cover around the ponds in 20 m perimeter (expressed as percentage) and the nearest pond distance (m). We considered that in the case of the temporary ponds these three pond variables are highly sensitive to environmental variations and have strong influence on the habitat quality of these ponds. We skipped other variables such as the vegetation cover in the ponds (due to low variability, the ponds being well vegetated), the presence of invertebrate predators (these were strongly associated with the more permanent water bodies) and the chemical properties of the ponds (we expected high variations within seasons and studies carried out in similar systems on amphibians shows no relationship between pond use and chemical variables, e.g. Plǎiaşu, Bǎncilǎ, Samoilǎ, Hartel & Cogǎlniceanu, 2012). In the following we will refer to those temporary ponds which were identified in the amphibian breeding period in 2009 and then in either 2018 and/or 2019 as “persisting temporary ponds (persisting ponds)” and to those which were not formed anymore in 2018 and 2019 as “lost temporary ponds (lost ponds)”.

Scrubs were represented in our study area by the Hawthorn (*Crataegus monogyna*), Blackthorn (*Prunus spinosa*) and the non-native and invasive Sea Buckthorn (*Hippophaë rhamnoides*) of which cover dramatically increased in the past 15-20 years. For this analysis we assessed the changes in scrub cover in (i) the whole study area for the period of 2003 (i.e. one year before the official expansion of the protected area), 2009 (the first pond and amphibian survey) and 2018 (the second pond and amphibian survey) and (ii) in a 20 m radius from the shoreline of each pond (for the year 2018). The scrub covers were estimated by vectorizing scrub patches. Google Earth Pro images. The images were georeferenced by using ArcGIS Desktop 10.3.1 software. As the spatial resolution of the images is satisfactory, the scale used for the vectorization process was 1:2000.

### 2.5 Understanding pasture management through interviews

We purposefully selected stakeholders as holders of knowledge and of long term genuine direct experience in relation to the management and recent history of the targeted pasture. By inquiring members of the local community about the existence of such persons, we were directed to three farmers. Most of the locals recognize these farmers as sources of knowledge about the management of the pasture and the history of the landscape. Their knowledge and experience was particularly salient especially in comparison to other community members who only very recently moved into the village and do not practice farming. Two farmers had 69 years (men) and one had 35 years (woman). The interviews with the farmers were conducted in the yard of their houses, placed close to the pasture. We further selected three ecologist professionals (two of them being conservationists) who also regularly visited the studied (case study) pasture during the past decade or earlier, for research or educational activities with students. The discussions were centered around the following three main topics: (i) Changes in the thorny scrub cover in the past decade (“Did you notice changes in scrub cover in the pasture in the past 10 years?”, “How you explain these changes?”, “How these changes influenced the quality of the pasture from the perspective of the local community?”). (ii) Weather conditions in the past decade (“How you appreciate the weather conditions in the past 10 years?”, “Where there any trends towards more or less rain?”, “How you appreciate the influence of the weather conditions on the pasture quality?”), (iii) Wetlands (“Did you notice any change in the wetland number from the pasture in the past decade?”, “How you explain these changes?”, “Is there any link between the scrub cover increase and the loss of small temporary wetlands?”. Ecologists were also queried about the possible ways in which scientists and local communities can work together for reviving the role of pasture for locals and their goals/aspirations/landscape preferences/landscape management priorities (“Do you see an opportunity in scientists working with local communities for reviving the role of pasture for the local communities? How?”). We took detailed notes while interviewing. Interviews were not recorded in order to encourage open responses. Given the specific topics addressed and the character of the local community (small number of people with the majority of persons recently settling there and not practicing farming) we are confident that the six experienced persons involved in the interviews provided a broad but accurate systems knowledge about the management and the state of the pasture. Their system knowledge serves to understand the current socio-economic changes and how to manage/navigate them for conserving the wetland biodiversity of the area, also valid for further/other restoration projects.

### 2.6 Data analysis

Below we will present the data analysis according of each objective (*O1-O4*, see above).

*O1*. We will present descriptive statistics only for the temporary ponds, because their persistence was threatened by land abandonment. We present the median and interquartile range (IQR) of the temporary pond variables for the three study years (2009, 2018 and 2019) and used Kruskal-Wallis test followed by the Nemenyi post-hoc test to compare these variables by using year (i.e. 2009, 2018, 2019) as grouping variable. We used Mann-Whitney U test to compare the variables describing the persisting ponds and the lost ponds (see above). In this analysis the pond type (i.e. persisting and lost) was the grouping variable. The five small sized permanent ponds were not considered in these analyses. We assessed the Euclidean distance between the nearest ponds (all ponds identified in a given year being considered, including the 5 permanent ponds) for every year (in m) to receive a simple metric pond isolation for the study area. We compared the yearly median pond distances with Kruskal-Wallis test.

*O2*. We used Generalized Linear Models (GLM) with binomial family to model the probability of temporary pond persistence according to the scrub cover (*log*) around the ponds (2018) and the pond area (*log*) in 2009 as predictor variables. The dependent variables were the persisting and lost ponds in the amphibian breeding season (see above for the definition of these two temporary pond types). We excluded the five permanent ponds which were mentioned historically (Nyárády, 1941), because the duration of these ponds does not depend on precipitation and is unlikely to be affected by grazing abandonment. Furthermore we excluded one lost temporary pond because it was drained.

*O3*. We used Non-Metric Multidimensional Scaling (NMDS) followed by the Analysis of Similarity (ANOSIM) to understand the differences in amphibian species assemblages in the lost (2009) and persisting (2009, 2018, 2019) as well as the permanent ponds. Presence-absence data of different amphibians in the ponds were used for this analysis. We compared the average amphibian species richness for the lost and persisting ponds with Analysis of Variance (ANOVA).

*O4*. Interviews were analyzed using open coding techniques to determine the themes mentioned and discussed by participants (Gibbs 2007). Then we created a short synthetic story about the management of the pasture. The results were shown to two persons (one interviewed and one new person with knowledge on the system) for validation.

### 2.7 Conceptualizing the social-ecological changes influencing amphibian breeding ponds

Based on our insights gathered from the ecological and social components of our results we developed an integrative causal loop dynamic on the management of pasture. The diagram resulting from this was then presented to one interviewed person and one new person knowing the pasture and corrected according to their suggestions. This system understanding will be presented in the Discussion section.

## 3. Results

### 3.1 The characteristics and dynamics of ponds (O1)

We identified 47 temporary ponds out of which 39 were not formed in 2018 and 2019 (i.e. lost ponds). Furthermore, we identified 11 new temporary ponds in the 2018 and 5 new temporary ponds in the 2019 survey. While the temporary pond area showed no significant variability in the three years (Table 1), the maximum depth was significantly higher in 2009 than in 2018 and 2019 (Table 1). Although the median area of the temporary ponds in the amphibian breeding season showed visible variability between the years, the difference was statistically not significant (Table 1). However, the upper quartile value of pond area shows a decreasing trend across the three years (Table 1). The maximum depth of the temporary ponds was smallest in 2018 and 2019 and highest in 2009 (Kruskal-Wallis H test and post-hoc test, P<0.05, Table 1). The disappearance of the temporary ponds resulted in an increase of the nearest pond distance from a median of 46 m (in 2009) to 68 m (in 2019), the differences being marginally significant (P = 0.07, Table 1). While the maximum depth of the lost and persisting ponds was the same (median value: 35), the upper quartile value of the depth was higher in the persisting ponds than that of the lost ponds (120 *vs* 50, Table 1). The area of the lost ponds was significantly smaller than the area of persisting ponds (Mann-Whitney U-test, P = 0.0002, Table 1). Figure 2 presents an example of temporary pond which was first identified in 2009 and it was formed in the amphibian breeding period in 2018 and 2019 as well.

**Table 1.** Descriptive statistics of pond variables for the three study years and according to the pond persistence. IQR=Interquartile-range.

| Variable name | 2009 | 2018 | 2019 | Kruskal-Wallis H test |
| --- | --- | --- | --- | --- |
|  | Median (IQR) | Median (IQR) | Median (IQR) |  |
| *Temporary ponds - between years* |  |  |  |  |
| Pond area | 150 (42-416) | 65 (24-177) | 85 (45-179) | P = 0.08 |
| Maximum depth | 35 (20-50) | 16 (12-23) | 16 (13-20) | P = 0.001 |
| Nearest pond distance | 46 (30-66) | 50 (36-114) | 68 (36-128) | P = 0.07 |
| *Persisting vs lost ponds* |  |  |  |  |
|  | Persisting ponds |  | Lost ponds | Mann-Whitney U test |
|  | Median (IQR) |  | Median (IQR) |  |
| Pond area | 110 (33-258) |  | 35 (20-50) | 0.0004 |
| Maximum depth | 35 (20-50) |  | 35 (20-120) | 0.64 |

### 3.2 Change in scrub cover and its relation to pond persistence (O2)

The overall scrub cover in the study area was 51 ha in 2003 and this increased to 102 ha in 2009 and then 151 ha in 2018 (Figure 3). Overall this means an increase of 296% for the period of 2003-2018 and 148% for the period of 2009-2018. The GLM showed that the probability of persistence of temporary ponds was significantly positively influenced by the pond area and significantly negatively influenced by the scrub cover around the ponds in 2018 (Table 2).

**Figure 3.**
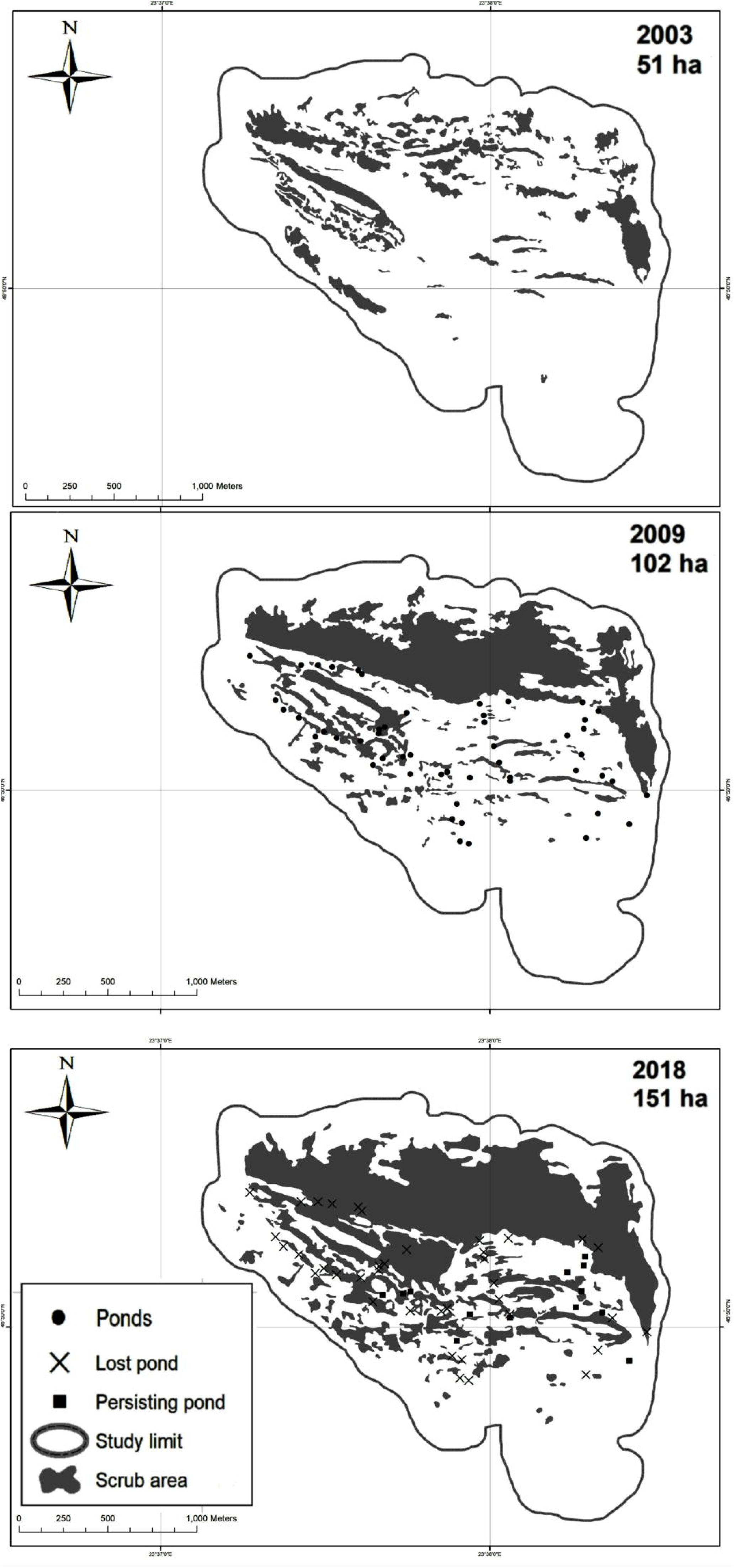
The overall change of scrub cover in the study area. The persisting and lost temporary ponds are also shown. No pond data are available for 2003.

**Table 2.** The relationship between the probability of temporary pond formation in the amphibian breeding and larval development period and the pond area and scrub cover around the temporary ponds.

| Variable | Estimate | SE | P |
| --- | --- | --- | --- |
| Intercept | -5.03 | 2.07 | 0.01 |
| Pond area (2009) | 0.79 | 0.36 | 0.02 |
| Scrub (2018) | -2.28 | 0.13 | 0.02 |

### 3.3 Amphibian use of ponds (O3)

During the three surveys, six amphibian species and a species complex were identified: *Triturus cristatus*, *Lissotriton vulgaris ampelensis*, *Bombina variegata*, *Rana dalmatina*, *Pelobates fuscus*, *Hyla arborea*, and a species complex *Pelophylax esculentus* and *P. ridibundus* (hereafter *P.* complex). *P. fuscus* was identified once in 2018. The occurrence in ponds sharply dropped during the years in *T. cristatus*, *L. vulgaris*, *B. variegata* and *H. arborea* (Figure 4) while *R. dalmatina* and the *P.* complex had stable pond use in 2018 and 2019 (Figure 4).

**Figure 4.**
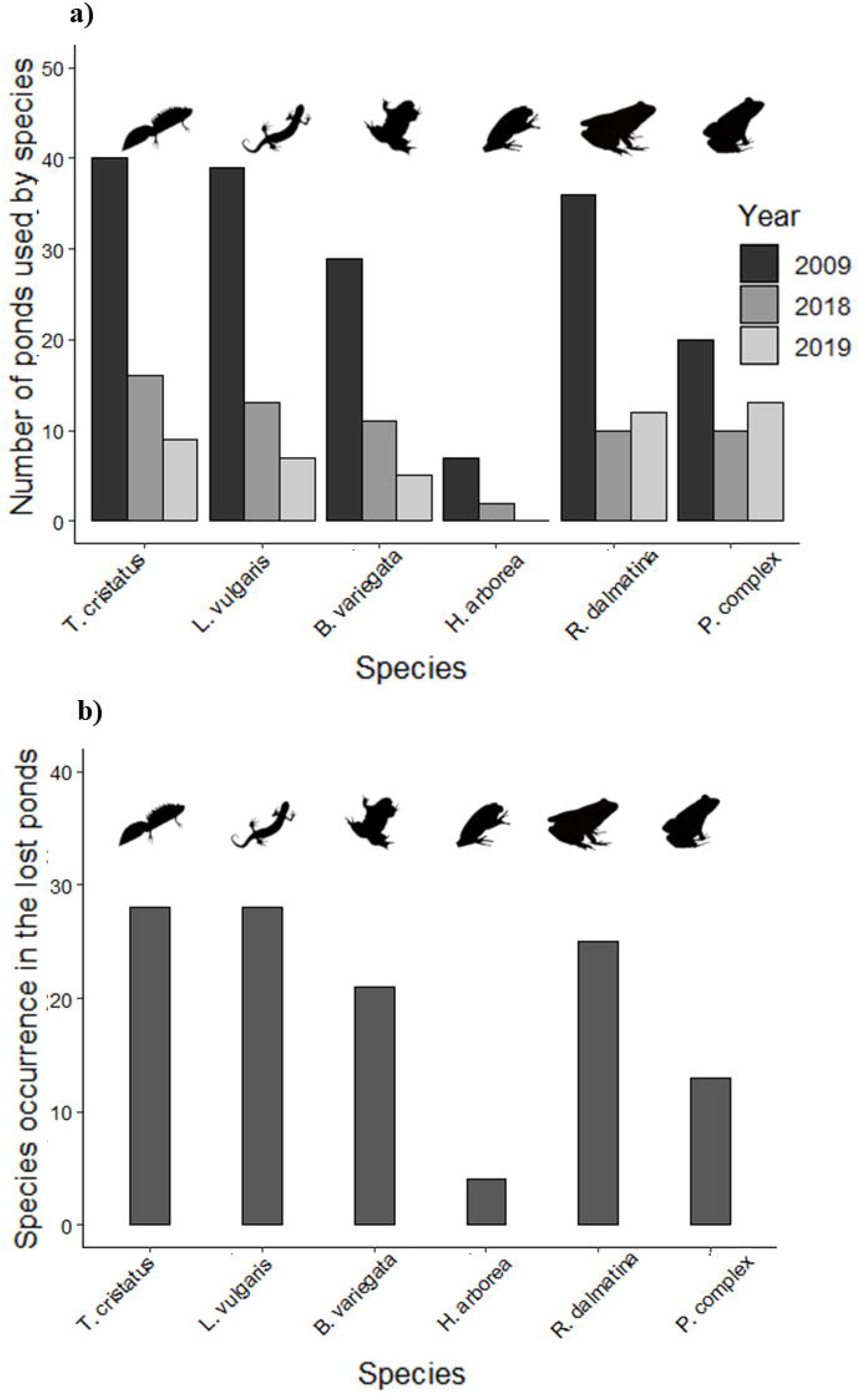
The presence of amphibians in the ponds in the study period (a) and the amphibian use of the lost temporary ponds (b).

Four species *T. cristatus, L. vulgaris, B. variegata* and *R. dalmatina* had over 20% occurrence in the lost ponds (Figure 4). The amphibian species richness was 2.92 [SD=1.02] for the lost ponds, 3.69 [1.18] for the persisting ponds in 2009, 2.90 [1.30] for the persisting ponds in 2018 and 3.00 [1.41] for the persisting ponds in 2019, these differences being not significant (ANOVA F_[3, 67]_=1.56, P=0.20). The NMDS and the subsequent ANOSIM analysis did not showed significant differences in the amphibian species assemblages in the lost and persisting ponds (P = 0.6 for ANOSIM statistic, other statistics not shown). The temporary ponds were dried till the end of May in 2018 and 2019 (Figure 2) having no detectable reproductive success.

### 3.4 Changes in grassland management as perceived by farmers and experts (O4)

All interviewees reported a dramatic increase in the scrub cover in the past decade or more and they linked the increased scrub cover to the decrease and complete abandonment of traditional pasture management and grazing. “*This was the pasture of the cattle and few buffalo, it was very clean, and you did not find a thorn on it. If you clear the thorns from the pasture so that the cattle have enough space to eat and walk, you will have also several butterflies around. It hurts my soul to see how the pasture changed.* (Interviewee 1, farmer).

“*When I studied the grassland for my master degree in the late 90’s I had big difficulties in finding a scrub where I can put my traps, the scrubs were so rare. Now we are lost in the scrubs*” (Interviewee 1, ecologist). The farmers related the increase of scrub cover to the decrease of pasture quality for livestock and also to the increased effort required to remove the dense scrub and restore the pasture quality in the pre-scrub encroachment period. One interviewee described in detail how the ill interpretation and enforcement of law amplified the abandonment of the pasture, triggered illegal grazing and then pasture abandonment and scrub encroachment in the pasture: “*Gradually, with the decrease in the number of livestock per capita, the interest on the grassland started to decrease and the scrubs were not cleared every year. The situation changed radically in 2004 when the reserve was extended by a Governmental Order (2151/2004) from 1.5 ha (see Law 5/2000) to 97 which included traditionally managed pasture. From an erroneous interpretation of the above mentioned Law (interpreting that protected area implicitly means the prohibition of any human actions) the Mayoralty of Cluj-Napoca ceased to rent the newly protected pasture surface to the locals. With this a paradox situation emerged: a decision that should have benefited the environment (i.e. the protected area extension) came to a situation that negatively affected both the local community and the ecosystem from the reserve. By prohibiting the traditional grazing without responsibilities was triggered. There were situations when locals brought their livestock during the night in the reserve to graze, sometimes even intensively. Being considered illegal activity, the grazing was slowly abandoned and this triggered the increase of thorny scrubs.*” (Interviewee 2, ecologist).

When asked about weather conditions and the wetlands from the pasture, the farmers and ecologists had distinct but complementary opinions. Farmers pointed towards the decrease of water levels in their fountains and the complete desiccation of a spring which normally did not dried in the past years. Furthermore, farmers mentioned about the existence of watering troughs for livestock and a natural pond with clean water in the 80`s. These all disappeared after the collapse of the communism due to destruction and lack of maintenance. The ecologists remarked that there is a decrease in the duration of temporary ponds as well as their number in the past years due to drought. The relation between scrub cover increase and loss of wetlands was not excluded by the stakeholders.

Ecologists see the collaboration between local farmers and conservationists as win-win situation for the local community and the pasture biodiversity, through educational and economic activities such as brands for local products. One ecologist highlighted that since 2014 when a conservation organization take the custody of the protected areas in the pasture, meetings were organized with members of the local communities to explore the possibilities to reinstall extensive grazing. Since 2018 extensive grazing is again possible for locals, which started to remove the scrubs in restricted pasture areas.

## 4. Discussion

We found that there is a dramatic increase of scrub cover in the study area and that the scrub cover negatively affects the formation of the temporary ponds in critical periods for amphibian reproduction and larval development. However, we did not find differences between the amphibian species assemblages nor the species richness between the lost- and persisting temporary ponds and the temporary and permanent ponds. The interviews with the local community provided a glimpse into the complex social, economic and institutional system interactions resulting in pasture abandonment and low motivation to restart traditional grazing.

### 4.1 Pond loss as a result of abandonment and their consequences on amphibians

In the studied 10-year period we documented the loss of 39 temporary ponds. The probability of temporary pond persistence across one decade in the breeding period of amphibians was negatively predicted by the increased scrub cover. Livestock (especially cattle and buffalo) grazing and trampling can prohibit the establishment of pond vegetation which otherwise would fill the ponds and/or increase their evapotranspiration (Boyce, Durtsche & Fugal, 2011; Pyke & Marty, 2005; Warren et al., 2007; Warren & Collins, 1994). Extensive grazing with livestock (cattle, buffalo) was related to the persistence of amphibian ponds in Europe and other continents (reviewed by Howell et al., 2019). The increase in scrub cover on pastures as a result of farming abandonment can result in loss of temporary ponds through the acceleration of desiccation (Boyce, Durtsche & Fugal, 2011; Ruiz, 2008). In turn, the accumulation of vegetation biomass and woody vegetation can decrease the quality of amphibian breeding habitats (Pyke and Marty, 2005; Skelly, Werner & Cortwright, 1999).

In patchy amphibian population systems (*sensu* Petranka & Holbrook, 2006) amphibians can assess the quality of aquatic habitats by within seasonal movements between ponds (Hartel, 2008). It is expected that the disappearance of the ponds and / or the decrease of their quality narrows the habitat diversity for several species, forcing amphibians to breed in suboptimal – i.e. sink- habitats (Pulliam, 2014) and decreasing the population and metapopulation viability in changing climate (Davis, Lohr & Roberts, 2018). The nearest pond distance in our study as a proxy of pond connectivity showed a slight increase with the pond loss, but the overall distances even with the loss of temporary ponds (Table 1) remained within the movement range of most species (i.e. less than 300 m) (Cayuela et al., 2014; Hartel, 2008). While the temporary character of the ponds may benefit amphibians because of the periodical removal of predators (Wellborn, Skelly & Werner, 1996), it was estimated that the duration which would maintain the suitability of the pond for self-sustaining local populations for species with long larval period would be one drying event in a decade (Oldham, Keeble, Swan & Jeffcote, 2000). In our study area none of the temporary ponds categorized by us as temporary would have such a long duration profile. We did not find significant difference between the amphibian species assemblages in the historically documented permanent ponds (Nyárády, 1941), and the persisting and lost temporary ponds (see Results). This suggests that the loss of the temporary ponds in the studied system does not affect the beta diversity of amphibians (characteristic to the whole pasture) albeit can reduce the alpha (i.e. pond specific) diversity. We assume that the persisting temporary and the permanent ponds act as refuges for amphibians, a situation which was described in other similar systems from Transylvania (Hartel, 2008; Hartel, Bǎncilǎ & Cogǎlniceanu, 2011).

### 4.2 The social aspects of amphibian pond conservation in traditional farming systems

In this study we also addressed the social dimensions of farmland abandonment (which has negative consequences on the formation of amphibian ponds, see above) in order to gather insights on the challenges and opportunities related to the conservation of temporary ponds in farming landscapes. Several studies highlighted the need to empower farmers for the conservation management of farmlands with high natural values (Molnár et al., 2015; Sandberg & Jakobsson, 2018; Snoo et al., 2013) or considering them in amphibian conservation initiatives in farming landscapes (Miró, O’Brien, Hall & Jehle, 2017; Semlitsch & Bodie, 2003). Conservation policy for traditional farming systems such as that addressed by us, should identify those socioeconomic, knowledge and management features which are shared by conservationists as well as the local community and build on these, in order to maintain environmentally friendly farming practices attractive (Fischer, Hartel & Kuemmerle, 2012). This is especially important because several protected species and habitats in the European Union (Halada, Evans, Romão & Petersen, 2011) and the world (Wright, Lake & Dolman, 2012) strongly depends on the continuation of extensive, wildlife friendly farming practices. Our interviews highlighted that farmers and ecologists agreed that maintaining open pasture surfaces with extensive cattle grazing is good for biodiversity while the excessive encroachment of scrub due to abandonment has negative consequences on pasture quality in terms of economy and habitats for different species (see Results). Similarly, Molnár et al. (2015) found several overlapping visions and objectives between herders and conservationists regarding the management of high biodiversity pastures of Hungary, including the maintenance of pasture surface by extensive grazing, maintenance of scattered woody vegetation across the pasture and the control of scrub encroachment. Furthermore, there was an agreement between herders and conservationists regarding the controlled maintenance of wetlands across the pasture, although for different purposes (Molnár et al., 2015). Our research also showed that the extension of the protected area into the managed pasture without the agreement of the local community and the complete prohibition of the grazing by the authorities resulted in the escalation of “illegal” grazing activities and did not stop occasional overgrazing. Our overall impression based on the interviews is that the interactions between the conservationists and the local communities in the early 2000`s does not allowed a careful mainstreaming of conservation laws into the local communities` value systems, knowledge types and farming practices. Difficulties around the interpretation and implementation of conservation policies in traditional farming systems from Transylvania were also highlighted from other regions (Mikulcak, Haider, Abson, Newig & Fischer, 2015). Figure 5 presents a causal loop diagram summarizing the social-ecological dynamics around pasture management and how this influenced the quality of aquatic habitats for amphibians. The pasture abandonment was driven by local socio-economic context (the decrease of profitability of traditional farming, strongly linked with the emigration of youth and the pasture overgrazing by locals and opportunistic outsiders, Figure 5), institutional context (the extension of protected area on managed pastures and the prohibition of grazing by the ill interpretation of conservation law, Figure 5) and the soil drought from the past decade (Figure 5). The abandonment of the pasture coupled with the decrease of pasture quality as the result of scrub encroachment as well as the strict application of nature conservation law were perceived as demotivating factors for reviving traditional farming in the pasture (see the reinforcing loop, R, in Figure 5). The overall abandonment of pasture and the scrub encroachment, with potential influence of weather conditions resulted in the loss of temporary ponds for amphibians (Figure 5). The system dynamic presented in the Figure 5 can be considered archetypical for the traditional management of communal pastures in Transylvania (Hanspach et al., 2014; Hartel et al., 2016; F. Mikulcak, Newig, Milcu, Hartel & Fischer, 2013; Sutcliffe et al., 2013) and elsewhere (Neudert, Salzer, Allahverdiyeva, Etzold & Beckmann, 2019) where the capacity of the local communities to self-organize in order to better navigate institutional challenges and opportunities were highlighted.

**Figure 5.**
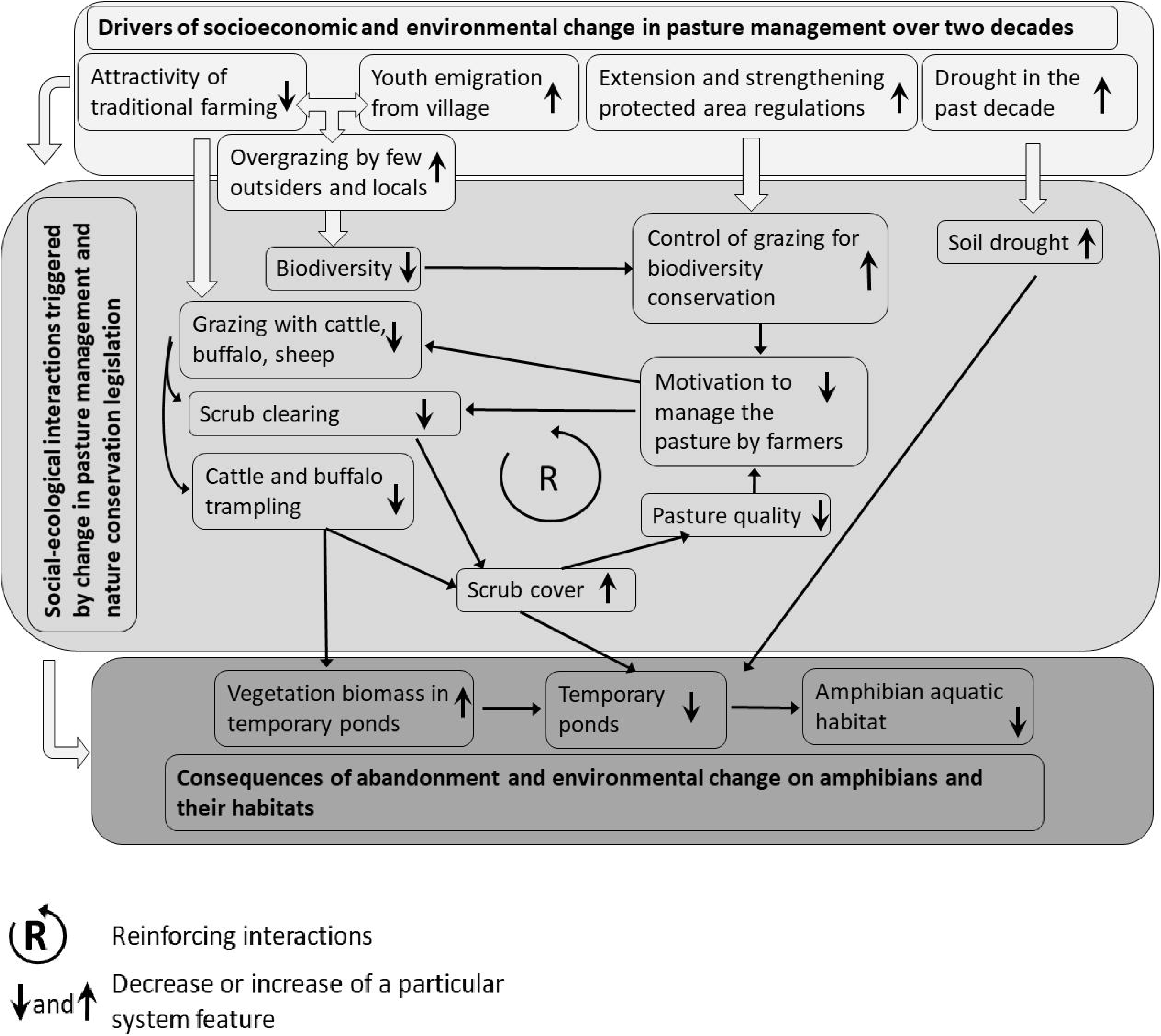
A system understanding on the social, economic and environmental drivers of temporary pond loss in pastures as a result of the abandonment of grazing.

### 4.3 Conclusions and conservation implications

Our study has two implications for the amphibian conservation management for regions which are similar to our study area. First, following the classical amphibian conservation paradigm, we suggest the prioritization of the remaining permanent and temporary ponds for amphibian conservation management strategies. Key elements of such a strategy are the manual control of excessive vegetation from selected temporary as well as the permanent ponds and controlling the scrubs in the vicinity of the temporary ponds. These actions, if carefully prioritized, are cost effective and can be implemented through volunteer and educational projects, to which the closely situated Cluj-Napoca city provides an excellent opportunity. Second, following a holistic, social-ecological perspective, we suggest the consideration of social-ecological system dynamics around pasture management in order to create social support for amphibian conservation initiatives at the level of local farmers. In this respect we identified a reinforcing loop around scrub encroachment, low pasture quality and the people`s demotivation to restart traditional grazing, despite the new attempts from a conservation NGO to revive traditional grazing practices. This reinforcing loop creates a social-ecological dynamic which is unfavorable for wetland conservation for amphibians. Furthermore, we suggest conservationists to identify management components which are favorable for amphibians and which are positively perceived by both farmers and conservationists (in our case the need for continuing traditional grazing with cattle and controlling scrubs) and join initiatives to address the attractivity and sustainability of these management actions for the local communities.

## Acknowledgements

Thanks to Vizauer Csaba, Petra Kulcsár, Zsombor Miholcsa, Edgár Papp, Brigitta-Ildikó Simon and István-Ervin Szegedi for helping during field surveys. EN was supported by a grant from Council of Harghita County and Association for Harghita County. The contribution of TH and CM was part of the project SusTaining AgriCultural ChAnge Through ecological engineering and Optimal use of natural resources (STACCATO), supported by a grant of the Romanian National Authority for Scientific Research and Innovation, CCCDI– UEFISCDI, project code ERA-FACCE-STACCATO-3..

## References

Howell, H. J., Mothes, C. C., Clements, S. L., Catania, S. V., Rothermel, B. B. & Searcy, C. A. (2019). Amphibian responses to livestock use of wetlands: new empirical data and a global review. Ecological Applications, in press, 0–2, https://doi.org/10.1002/eap.1976

Alford, R. & Richards, S. (1999). Global Amphibian Declines: A Problem. Annual Review of Ecology, Evolution, and Systematics, 30, 133–165. https://doi.org/10.1146/annurev.ecolsys.30.1.133

Bergmeier, E., Petermann, J. & Schröder, E. (2010). Geobotanical survey of wood-pasture habitats in Europe: diversity, threats and conservation. Biodiversity and Conservation, 19, 2995–3014. https://doi.org/10.1007/s10531-010-9872-3

Bielz, E.A. (1888). Die Fauna der Wierbeltiere Siebenbürgens nach ihrem jetzigen Bestande, Verhandlungen und Mittheilungen des Siebenbürgischen Vereins für Naturwissenschaften in Hermannstadt, 39, 15–120

Boix, D., Biggs, J., Hull, A.P., Kalettka, T. & Oertli, B. (2012). Pond research and management in Europe: “Small is Beautiful.”, Hydrobiologia, 689, 1–9. https://doi.org/10.1007/s10750-012-1015-2

Ruhí, A., San Sebastian, O., Feo, C., Franch, M., Gascón, S., Richter-Boix, A., Boix, D. & Llorente, G. (2012). Man-made Mediterranean temporary ponds as a tool for amphibian conservation, Annales de Limnologie, 48, 81–93. https://doi.org/10.1051/limn/2011059

Mikulcak, F., Haider, J. L., Abson, D. J., Newig, J., & Fischer, J. (2015). Land Use Policy Applying a capitals approach to understand rural development traps: A case study from post-socialist Romania. Land Use Policy, 43, 248–258. https://doi.org/10.1016/j.landusepol.2014.10.024

Boyce, R.L., Durtsche, R.D. & Fugal, S.L. (2011). Impact of the invasive shrub *Lonicera maackii* on stand transpiration and ecosystem hydrology in a wetland forest. Biological Invasions, 1–20. https://doi.org/10.1007/s10530-011-0108-6

Miró, A., O’Brien, D., Hall, J. & Jehle, R. (2017). Habitat requirements and conservation needs of peripheral populations: the case of the great crested newt (*Triturus cristatus*) in the Scottish Highlands. Hydrobiologia, 792(1), 169–181. https://doi.org/10.1007/s10750-016-3053-7

Buckley, J., Beebee, T.J.C. & Schmidt, B.R. (2012). Monitoring amphibian declines: population trends of an endangered species over 20 years in Britain. Animal Conservation, 17, 27–34, https://doi.org/10.1111/acv.12052

Buono, V., Bissattini, A.M. & Vignoli, L. (2019). Can a cow save a newt? The role of cattle drinking troughs in amphibian conservation, Aquatic Conservation: Marine and Freshwater Ecosystems, 29, 964–975. https://doi.org/10.1002/aqc.3126

Calhoun, A.J.K., Mushet, D.M., Bell, K.P., Boix, D., Fitzsimons, J.A. & Isselin-Nondedeu, F. (2017). Temporary wetlands: challenges and solutions to conserving a ‘disappearing’ ecosystem. Biological Conservation, 211, 3–11. https://doi.org/10.1016/j.biocon.2016.11.024

Cayuela, H., Besnard, A., Bonnaire, E., Perret, H., Rivoalen, J. & Miaud, C. (2014). To breed or not to breed: past reproductive status and environmental cues drive current breeding decisions in a long C lived amphibian. Oecologia, 176, 107–116, https://doi.org/10.1007/s00442-014-3003-x

Cogǎlniceanu, D., Bǎncilǎ, R., Plǎiaşu, R., Samoilǎ, C. & Hartel, T. (2012). Aquatic habitat use by amphibians with specific reference to *Rana temporaria* at high elevations (Retezat Mountains National Park, Romania). Annales de Limnologie, 48, 355–362, https://doi.org/10.1051/limn/2012026

Curado, N., Hartel, T. & Arntzen, J.W. (2011). Amphibian pond loss as a function of landscape change - A case study over three decades in an agricultural area of northern France. Biological Conservation, 144, 1610–1618, https://doi.org/10.1016/j.biocon.2011.02.011

Davis, R.A., Lohr, C.A. & Roberts, J.D. (2018). Frog survival and population viability in an agricultural landscape with a drying climate. Population Ecology, 61, 102–112, https://doi.org/10.1002/1438-390X.1001

Demeter, L. & Hartel, T. (2007). A rapid survey of large branchiopods in Romania. Annales de Limnologie, 43, 101–105, https://doi.org/10.1051/limn/2007016

Ferreira, M. & Beja, P. (2013). Mediterranean amphibians and the loss of temporary ponds: Are there alternative breeding habitats? Biological Conservation, 165, 179–186. https://doi.org/10.1016/j.biocon.2013.05.029

Fischer, J., Hartel, T. & Kuemmerle, T. (2012). Conservation policy in traditional farming landscapes. Conservation Letters, 5, 167–175, https://doi.org/10.1111/j.1755-263X.2012.00227.x

Flitcroft, R., Boon, P. J., Cooperman, M. S., Harrison, I. J., & Bignoli, D. J. (2019). Theory and practice to conserve freshwater biodiversity in the Anthropocene, Aquatic Conservation: Marine and Freshwater Ecosystems, 1013–1021. https://doi.org/10.1002/aqc.3187

Halada, L., Evans, D., Romão, C. & Petersen, J.-E. (2011). Which habitats of European importance depend on agricultural practices? Biodiversity and Conservation, 20, 2365–2378. https://doi.org/10.1007/s10531-011-9989-z

Hanspach, J., Hartel, T., Milcu, A.I., Mikulcak, F., Dorresteijn, I., Loos, J., … Fischer, J. (2014). A holistic approach to studying social-ecological systems and its application to Southern Transylvania. Ecology and Society, 19. https://doi.org/10.5751/ES-06915-190432

Hanspach, J., Loos, J., Dorresteijn, I., Abson, D.J. & Fischer, J. (2016). Characterizing social– ecological units to inform biodiversity conservation in cultural landscapes. Diversity and Distributions, 22, 853–864. https://doi.org/10.1111/ddi.12449

Hartel, T. (2008a). Movement activity in a *Bombina variegata* population from a deciduous forested landscape. North-Western Journal of Zoology, 4, 79–90

Hartel, T. (2008b). Long-term within pond variation of egg deposition sites in the agile frog, *Rana dalmatina*. Biologia (Bratislava), 63, 439–443, https://doi.org/10.2478/s11756-008-0060-9

Hartel, T., Bǎncilǎ, R. & Cogǎlniceanu, D. (2011). Spatial and temporal variability of aquatic habitat use by amphibians in a hydrologically modified landscape. Freshwater Biology, 56, 2288–2298, https://doi.org/10.1111/j.1365-2427.2011.02655.x

Hartel, T., Fagerholm, N., Torralba, M., Balázsi, Á. & Plieninger, T. (2018). Forum: Social-Ecological System Archetypes for European Rangelands. Rangeland Ecology and Management, 71, 536–544, https://doi.org/10.1016/j.rama.2018.03.006

Hartel, T., Olga Réti, K., Craioveanu, C., Gallé, R., Popa, R., Ioniţă, A., Demeter, L., Rákosy, L. & Czúcz, B. (2016). Rural social–ecological systems navigating institutional transitions: case study from Transylvania (Romania). Ecosystem Health and Sustainability, 2, https://doi.org/10.1002/ehs2.1206

Hartel, T. & von Wehrden, H. (2013). Farmed areas predict the distribution of amphibian ponds in a traditional rural landscape. PLoS One 8, e63649. https://doi.org/10.1371/journal.pone.0063649

Klein Tank, A.M.G., Wijngaard, J. B., Können, G.P., Böhm, R., Demarée, G., Gocheva, A., … Petrovic, P. (2002). Daily dataset of 20th-century surface air temperature and precipitation series for the European Climate Assessment. International Journal of Climatology, 22, 1441–1453. https://doi.org/10.1002/joc.773 Data and metadata available at http://www.ecad.eu

Merilä, J., Laurila, A., Pahkala, M., Räsänen, K. & Timenes, A. (2010). Adaptive phenotypic plasticity in timing of metamorphosis in the common frog *Rana temporaria*, Ecoscience, 7, 18–24. https://doi.org/10.1080/11956860.2000.11682566

Mikulcak, F., Newig, J., Milcu, A.I., Hartel, T. & Fischer, J. (2013). Integrating rural development and biodiversity conservation in Central Romania. Environmental Conservation, 40. https://doi.org/10.1017/S0376892912000392

Molnár, Z., Kis, J., Vadász, C., Papp, L., Sándor, I., Béres, S., Sinka, G. & Varga, A. (2015). Common and conflicting objectives and practices of herders and conservation managers: the need for a conservation herder. Ecosystem Health and Sustainability, 1, 1–20. https://doi.org/10.1002/ehs2.1215

Neudert, R., Salzer, A., Allahverdiyeva, N., Etzold, J. & Beckmann, V. (2019). Archetypes of common village pasture problems in the South Caucasus: insights from comparative case studies in Georgia and Azerbaijan. Ecology and Society, 24, 5.

Newman, R.A. (1992). Adaptive Plasticity in Amphibian Metamorphosis. What type of phenotypic variation is adaptive, and what are the costs of such plasticity?. Bioscience, 42, 671–678.

Nyárády, E.Gy. (1941). A kolozsvári Szénafüvek suvadásos területeiről. Földrajzi közlemények, Budapest, 40–53.

Oldham, R.S., Keeble, J., Swan, M.J.S. & Jeffcote, M. (2000). Evaluating the habitat suitability of the great crested newt (*Triturus cristatus*). Herpetological Journal, 10, 143–155.

Petranka, J.W. & Holbrook, C.T. (2006). Wetland Restoration for Amphibians: Should Local Sites Be Designed to Support Metapopulations or Patchy Populations? Restoration Ecology, 14, 404–411.

Plǎiaşu, R., Bǎncilǎ, R., Samoilǎ, C., Hartel, T. & Cogǎlniceanu, D. (2012). Waterbody availability and use by amphibian communities in a rural landscape. Herpetological Journal, 22.

Plieninger, T., Hartel, T., Martín-López, B., Beaufoy, G., Bergmeier, E., Kirby, … Van Uytvanck, J. (2015). Wood-pastures of Europe: Geographic coverage, social-ecological values, conservation management, and policy implications. Biological Conservation, 190, 70–79. https://doi.org/10.1016/j.biocon.2015.05.014

Pulliam, R. (2014). Sources, sinks and population regulation. The American Naturalist, 132, 652–661.

Pyke, C.R. & Marty, J. (2005). Cattle Grazing Mediates Climate Change Impacts on Ephemeral Wetlands, Conservation Biology, 1619–1625. https://doi.org/10.1111/j.1523-1739.2005.00233.x

Richter-boix, A., Tejedo, M. & Rezende, E.L. (2011). Evolution and plasticity of anuran larval development in response to desiccation. A comparative analysis. Ecology and Evolution, 15–25. https://doi.org/10.1002/ece3.2

Ruiz, E. (2008). Management of Natura 2000 habitats - Mediterranean temporary ponds. Technical Report, European Commission.

Ruprecht, E. (2017). Secondary succession in old-fields in the Transylvanian Lowland (Romania). Preslia 77, 145–157.

Sandberg, M. & Jakobsson, S. (2018). Trees are all around us: Farmers’ management of wood pastures in the light of a controversial policy. Journal of Environmental Management, 212, 228–235. https://doi.org/10.1016/j.jenvman.2018.02.004

Scheffer, M., Van Geest, G. J., Zimmer, K., Jeppesen, Søndergaard, M., E., Butler, M.G., … Meester, L. De. (2006). Small habitat size and isolation can promote species richness: second-order effects on biodiversity in shallow lakes and ponds, OIKOS, 227–231.

Semlitsch, R.D. (2000). Principles for management of aquatic-breeding amphibians, Journal of Wildlife Management, 64, 616–631.

Semlitsch, R.D. & Bodie, J.R. (1998). Are Small, Isolated Wetlands Expendable? Conservation Biology, 12, 1129–1133.

Semlitsch, R.D. & Bodie, J.R. (2003). Biological Criteria for Buffer Zones around Wetlands and Riparian Habitats for Amphibians and Reptiles, Conservation Biology, 17, 1219–1228.

Semlitsch, R. D., Peterman, W. E., Anderson, T. L., Drake, D. L. & Ousterhout, B. H. (2015). Intermediate Pond Sizes Contain the Highest Density, Richness, and Diversity of Pond-Breeding Amphibians, PloS One, 1–20. https://doi.org/10.1371/journal.pone.0123055

Skelly, D.K., Werner, E.E. & Cortwright, S.A. (1999). Long term distributional dynamics of a Michigan amphibian assemblage. Ecology, 80, 2326–2337.

Snoo, G.R. De, Herzon, I., Staats, H., Burton, R.J.F., Schindler, S., Dijk, … Musters, C.J.M.. (2013). Toward effective nature conservation on farmland: making farmers matter, Conservation Letters, 6, 66–72. https://doi.org/10.1111/j.1755-263X.2012.00296.x

Sos, T. & Hegyeli, Zs. (2014). Characteristic morphotype distribution predicts the extended range of the “Transylvanian” smooth newt, *Lissotriton vulgaris ampelensis* Fuhn 1951, in Romania. North-Western Journal of Zoology, 11, 34–40

Sutcliffe, L., Paulini, I., Jones, G., Marggraf, R. & Page, N. (2013). Pastoral commons use in Romania and the role of the Common Agricultural Policy. International Journal of the Commons, 7, 58–72.

Tanadini, M., Schmidt, B.R., Meier, P., Pellet, J. & Perrin, N. (2012). Maintenance of biodiversity in vineyard-dominated landscapes: a case study on larval salamanders, Animal Conservation, 15, 136–141. https://doi.org/10.1111/j.1469-1795.2011.00492.x

Ustaoglu, E. & Collier, M.J. (2018). Farmland abandonment in Europe: an overview of drivers, consequences, and assessment of the sustainability implications. Environmental Reviews, 26, 1–21. https://doi.org/10.1139/er-2018-0001

Warren, S.D. & Collins, F. (1994). Relationship of Endangered Amphibians to Landscape Disturbance, Journal of Wildlife Management, 72, 738–744, https://doi.org/10.2193/2007-160

Warren, S.D., Holbrook, S.W., Dale, D.A., Whelan, N.L., Elyn, M., Grimm, W. & Jentsch, A. (2007). Biodiversity and the Heterogeneous Disturbance Regime on Military Training Lands, European Journal of Entomology, 15, 606–612. https://doi.org/10.14411/eje.2015.099

Wellborn, G.A., Skelly, D.K. & Werner, E.E. (1996). MECHANISMS CREATING COMMUNITY STRUCTURE ACROSS A FRESHWATER HABITAT GRADIENT. Annual Review of Ecology, Evolution, and Systematics, 27, 337–363

Wells, K.D. (1977). The social behaviour of anuran amphibians. Animal Behaviour, 25, 666–693. https://doi.org/10.1016/0003-3472(77)90118-X

Winter, T.C. (2000). The vulnerability of wetlands to climate change: a hydrologic landscape perspective. Journal of the American Water Resources Association, 36, 305–311.

Wright, H.L., Lake, I.R. & Dolman, P.M. (2012). Agriculture-a key element for conservation in the developing world. Conservation Letters, 5, 11–19. https://doi.org/10.1111/j.1755-263X.2011.00208.x

